# Quantifying vulnerability to plant invasion across global ecosystems

**DOI:** 10.1101/2024.02.21.581382

**Authors:** William G. Pfadenhauer, Bethany A. Bradley

## Abstract

The widely-referenced “tens rule” in invasion ecology suggests that 10% of established, non-native species will become invasive. However, the accuracy of this estimate has been questioned, as the original analysis focused on small groups of plant species in Great Britain and Australia. Using a novel database of 9,501 established and 2,924 invasive plants, we provide a comprehensive evaluation of plant invasion rates and the first empirical analysis of how the tens rule varies across climate zones and spatial scales. We found that invasion rates ranged from 17% at the country scale to 25% at the continental scale. Tropical communities are often considered to be resistant to invasion, however our results showed significantly higher invasion rates in the tropics and especially on tropical islands, suggesting unexpectedly high vulnerability of these species-rich ecosystems. Our analysis provides improved, environment-specific estimates of invasion rates which are often twice as high as previous expectations. We recommend that practitioners reject the tens rule for plants and adjust future management practices to reflect these updated estimates of invasion risk.

**Open Research Statement:** The data and code that support the findings of this study are openly available on GitHub at https://github.com/wpfadenhauer/Global-Invasion-Rates

## Introduction

Non-native invasive species are one of the principal drivers of biodiversity loss (Vitousek et al. 1997, Wilcove et al. 1998, Mollot et al. 2017, Nic Lughadha et al. 2020), have disrupted nutrient cycling and fire regimes (Brooks et al. 2004, Vilà et al. 2011), and are responsible for billions of dollars in economic losses each year (Pimentel et al. 2005). Predicting the percentage of non-native species that will establish self-sustaining populations when introduced to new regions, and the percentage of these established species that will invade (and have significant negative impacts) is critical for risk assessment and preventative management. The most common estimate for these percentages is the “tens rule,” which posits that 10% of introduced species will establish and 10% of these established species will become invasive (Williamson and Fitter 1996). The tens rule is widely cited (Enders et al. 2018) and commonly interpreted as evidence that the vast majority of introduced and established species will not become invasive and therefore risk of harm is low (Jarić and Cvijanović 2012). This ubiquitous interpretation assumes that data from Great Britain and Australia (Williamson and Fitter 1996) are an accurate representation of all global ecosystems.

Subsequent studies of the tens rule have typically focused on the first step: the transition from non-native introduction to establishment of self-sustaining naturalized populations. The second transition - from establishment to invasion - is essential for understanding risk of invasion and impact, but remains vastly understudied. Of the 21 papers that examined plants in a 2018 meta-analysis of the tens rule, only six examined the proportion of species making transition from establishment to invasion (hereafter “invasion rate”; Jeschke and Pyšek 2018). Of these, two were part of the original study (Williamson 1993, Williamson and Fitter 1996) and two studies replicated assessments of invasion rates in Australia (Scott and Panetta 1993, Virtue et al. 2004). These few studies in Great Britain and Australia are not representative of global ecosystems and have themselves shown inconsistent support for the tens rule, suggesting that a broader, global analysis is needed to understand this core issue in invasion ecology.

The shortcomings of past estimates of plant invasion rates can be partially attributed to an overall lack of consistent information about invasive plants at a global scale. For example, many countries do not regulate or document invasive plants (Early et al. 2016) and those that do show pronounced inconsistencies across internal political borders (e.g., Beaury et al. 2021). Moreover, different sources often use incompatible definitions of important terms which has made compiling disparate databases challenging. For example, some sources conflate establishment with invasion (e.g., Randall 2017), while others have differing standards for what qualifies a species as a “pest” (Jarić and Cvijanović 2012, Jeschke and Pyšek 2018). These issues have contributed to spatial biases in measurements of invasion rates. Fortunately, the recent development of global databases with consistent and standardized records of both established (van Kleunen et al. 2019) and invasive (Laginhas and Bradley 2021, CAB International 2022, Global Register of Introduced and Invasive Species 2022) plant species have created the opportunity for a new, more robust global assessment of invasion rates.

Inconsistent estimates of invasion rates may also have been caused by varying spatial scales of analyses. Larger study areas (e.g., continents) generally contain more habitat heterogeneity than smaller study areas (e.g., states, islands), potentially creating opportunities for invasion for a wider group of species, and ultimately leading to higher invasion rates. Although scale-dependence of invasion rates has been proposed (Jeschke and Pyšek 2018), it has yet to be demonstrated with empirical evidence.

Invasion rates are also likely to vary with geography and climate. The “island susceptibility hypothesis” suggests that invasion rates on islands should be higher than those on mainlands due to smaller species pools and the consequences of past colonization and intra-island evolution (Elton 1958, Simberloff 1995, Jeschke et al. 2018). Additionally, invasion rates in tropical climates have previously been hypothesized to be lower than those in other climates due to invasion resistance conferred from high native species richness (Elton 1958, Holdgate 1986, Lonsdale 1999). Both hypotheses are well-known (Enders et al. 2018) but have received mixed empirical support (Jeschke et al. 2018). This is particularly true for the assumption that tropical areas are invasion resistant, which is chronically under-studied (Chong et al. 2021). Therefore, a macroscale examination with newly-available, globally-integrated data would help reconcile the conflicting evidence that exists for these two hypotheses.

Here, we compiled a global database of established and invasive plant species which we used to calculate invasion rates. We used these invasion rates to determine: 1) whether 10% is an accurate estimate of global invasion rates for plants, 2) the influence that spatial scale might have in estimating invasion rates, and 3) which global environments are most susceptible to plant invasions. This study provides the first integrated assessment of how plant invasion rates vary across recipient systems at a global scale.

## Methods

### Lists of established and invasive species

We compiled a list of all plant species documented as established and/or invasive in one or more regions of the world. We identified a species as ‘established’ (having been introduced and having one or more self-sustaining populations outside the native distribution) if it was listed as “naturalized” in the most recent version of the Global Naturalized Alien Flora (GloNAF) database (van Kleunen et al. 2019). We identified a species as ‘invasive’ (established and spreading or having negative impacts) using the Global Plant Invaders database (Laginhas and Bradley 2021). We only defined species as invasive if they were listed as “Invasive explicitly defined” or “Invasive implicitly defined” in the Global Plant Invaders database, which provided higher confidence in the species’ invasive status. We subsequently added any additional plant species that we discovered through data collection as having documented negative impacts in at least one location (from CABI, GRIIS, and Zhang et al.; Zhang et al. 2021, CAB International 2022, Global Register of Introduced and Invasive Species 2022). Since inconsistent applications of the term “invasive” have previously been an obstacle to analyzing invasion trends in the past, we specifically chose sources of invasive plant distributions that used compatible methods for defining invasiveness. We combined these lists of established and invasive species and resolved any taxonomic inconsistencies using the ‘TNRS’ package in R (with source = “WCVP” and accuracy = 0.9). This created a list of 12,929 candidate species.

### Compiling native, established, and invaded regions

For each of the species in our list, we compiled its native distribution, established distribution, and invaded distribution (Table S1). These distributions were collected at the finest resolution reported by each of the online sources, which often coincided with political boundaries. We webscraped data using the R package ‘rvest’. For species with CABI ISC datasheets, we webscraped the “Distribution Tables,” specifically the locations listed as “Native” in the “Origin” column and the locations listed as “Invasive” in the “Invasive” column (CAB International 2022). For species listed in Plants of the World Online, we webscraped all locations under the “Native to” heading (Royal Botanic Gardens, Kew 2022). We were unable to find established or invaded regions for 504 species on our candidate list and thus excluded them from the analysis. This resulted in a final list of 12,425 species.

For each of the 12,425 species, we recorded whether it was native, established, or invasive within a given geographic region at four spatial scales, corresponding to L1 through L4 (Fig. 1) of the World Geographical Scheme for Recording Plant Distributions (WGSRPD; Brummitt 2001). We converted plant records at finer spatial scales to their corresponding coarser scales for completeness. For example, species reported as invasive in the state of Massachusetts, U.S.A. (an L3 region) were also recorded as invasive in the Northeastern U.S.A. (the corresponding L2 region) and Northern America (the corresponding L1 region). Because some geographical information was only reported at the country or continental level, our spatial database includes the most information at the L1 (continental) scale and the least information at the L4 (province/state) scale.

**Fig. 1.**
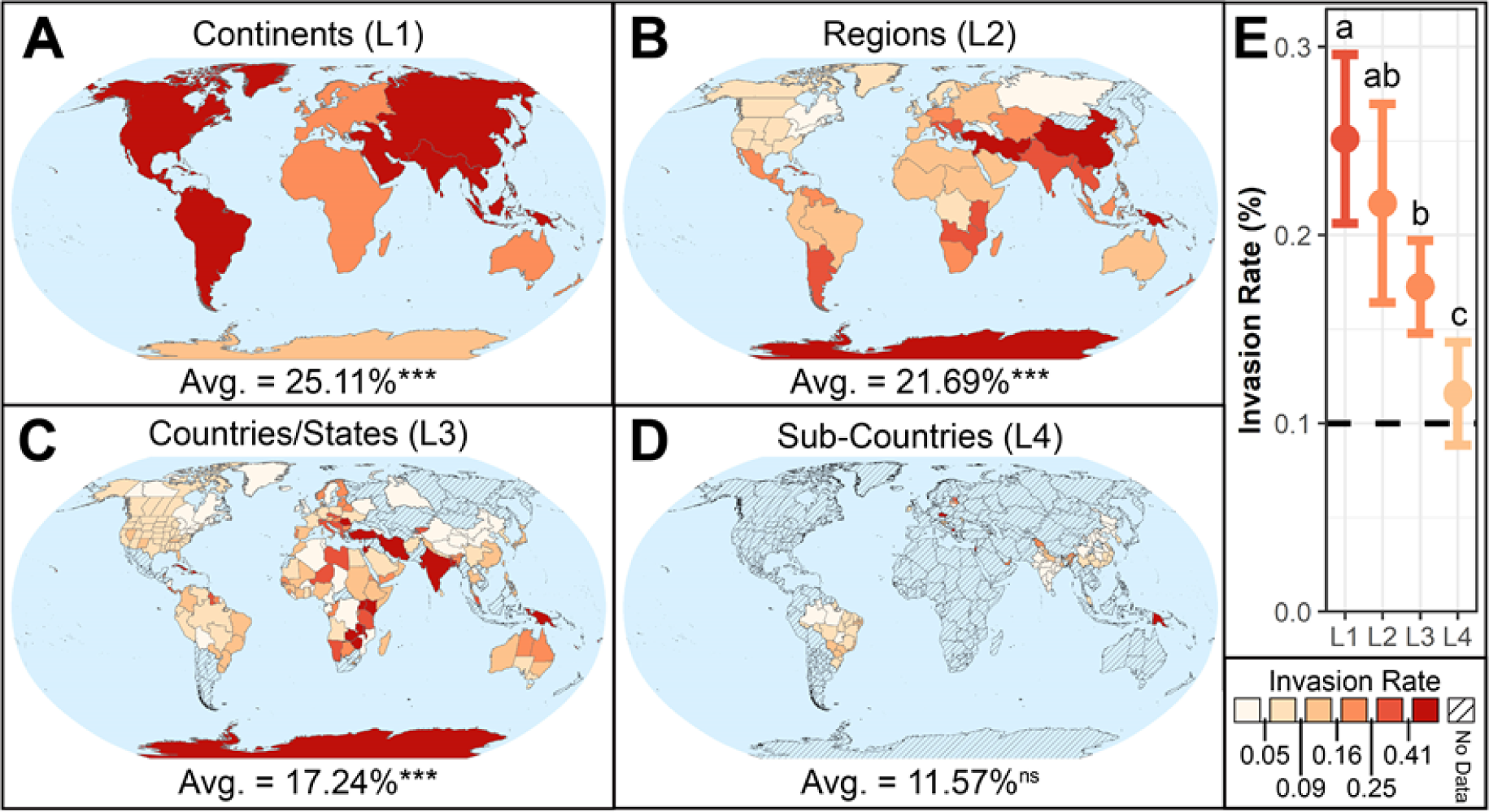
Invasion rates vary by scale, but at most practical scales they are significantly higher than predicted by the tens rule. Invasion rates of WGSRPD regions at the continental (L1) scale (A), regional (L2) scale (B), country/state (L3) scale (C), and sub-country (L4) scale (D). Superscripts denote whether the average at that scale is significantly different than 10% (^ns^not significant, ***p-value ≤ 0.001). (E): Means and 95% confidence intervals for the invasion rates calculated at each of the four scales. The dashed line represents the tens rule. Lowercase letters (a, b, c) indicate significant differences across scales (p-value ≤ 0.05). T-values, degrees of freedom, and p-values are provided in Table S2.

### Assigning island labels

We categorized all regions as either islands or mainlands to determine whether islands had higher susceptibility to invasion. We assigned island labels to the finest-scale WGSRPD region in each location. In some locations (like the continental United States) L3 is the finest classification used, whereas in other locations (like eastern Canada) L4 classifications are available. To distinguish islands and mainlands, we used the island labels provided in GloNAF (van Kleunen et al. 2019) and then manually assigned labels to any regions that were missing (n = 75 islands and 91 mainlands). Species without L3 or L4 destination data were excluded from the island analyses (n = 73 species).

### Assigning climate labels

Since it has long been suggested that different climates vary in their susceptibility to invasion, we also assigned climates to the finest scale WGSRPD region in each location (Holdgate 1986, Lonsdale 1999, van Kleunen et al. 2015). To distinguish between climates, we assigned to each region the Köppen climate that was most common within its boundaries according to Kottek et al. (2006). For example, the state of Massachusetts, U.S.A. encompasses four different climates (Cfa, Cfb, Dfa, and Dfb), with the largest area classified as Dfb (continental climate with no dry season and warm summers), thus we considered species native to Massachusetts as having a source climate of Dfb, and species established or invasive in Massachusetts as having a recipient climate of Dfb. Throughout the manuscript, we refer to major climates as defined by the Köppen Climate Classification (A - Tropical, B - Arid, C - Temperate, D - Continental, and E - Polar; Figure S2). Species without L3 or L4 destination data were excluded from climate analyses.

### Calculating invasion rates

To determine the validity of the tens rule across global ecosystems, we calculated the percentage of established plants in each region that are invasive. We intentionally used invasion rates instead of invasive plant richness because it enabled a direct comparison with the second transition included in the tens rule, and because it helps to correct for spatial biases in invasive plant distribution data. Sampling efforts for non-native plants are globally inconsistent, but analyses produced by GloNAF and the Global Plant Invaders database (our main sources of established and invasive plant species) suggest that areas which are under-sampled for established and invasive plants largely overlap (van Kleunen et al. 2019, Laginhas et al. 2022). Thus, dividing the number of invasive species by the number of established species helps to account for these spatial biases when analyzing global invasion patterns.

### Comparing invasion rates across scales

We averaged invasion rates across regions at all four WGSRPD levels. We excluded species that were native to the regions where they were listed as established/invasive because the tens rule typically focuses on non-native species introductions. Using two-sided t-tests and alpha values of 0.05, we tested for significant differences between means at each of the four average WGSRPD invasion rates, and also between each average invasion rate and the expected value of 10%.

### Island susceptibility analyses

To determine whether islands are more susceptible to invasions than mainlands (as is expected by the island susceptibility hypothesis), we compared invasion rates for islands and mainlands. We aggregated invasion rates at the global scale and we averaged invasion rates across regions at the L3 scale to account for scale-dependence. We chose L3 instead of L4 for our finest scale because L3 includes nearly twice as many species (12,352 vs. 6,816) and because invasive plant species are more commonly regulated at the L3 scale than the L4 scale. Since it is possible for species that are native to islands to be established and/or invasive in other islands, we retained species that were native to the same category of region (i.e., “island” or “mainland”) where they invaded. We compared island and mainland invasion rates at the global scale using a one-sided t-test with an alpha value of 0.05. Since data aggregated at the L3 scale consisted of invasion rates for 51 island regions and 179 mainland regions, we calculated average invasion rates for islands and mainlands weighted by the total number of species in each individual region. We compared these weighted means using a weighted two-sample, one-sided t-test, with the alternate hypothesis that islands had higher invasion rates than mainlands.

### Climate comparisons

To account for potential differences in island susceptibility caused by differences in climate susceptibility, we calculated the invasion rates in each major Köppen climate. We also calculated invasion rates for the 31 subclimates in the Köppen classification system, although 9 of these categories lacked records of invasive species (Csc, Cwc, Dfd, Dsa, Dsb, Dsc, Dsd, Dwc, and Dwd), so we only included results for the 5 major climates. Since it is possible for species to be established and/or invasive in non-native regions that have the same major climate as their native regions, we retained species that were native to the same climate(s) where they had invaded. We compared percentages across the major climates using a pairwise proportion test (‘pairwise.prop.test’ function in the R package ‘stats’) and an alpha value of 0.05 with the Bonferroni correction. We ran these comparisons three times: once with all applicable regions, once with islands only, and once with mainlands only.

In addition to calculating incoming invasion rates for each climate, we also calculated the invasion rates “donated” by each climate to determine if some climates were more likely to be sources of invasive plants (Fig. S1).

## Results

We identified 12,425 plant species that had become established in at least one location. We found native distributions for 12,170 of these established species, and we found evidence of invasion in at least one location for 2,924 of the established species. Therefore, at a global scale, the percentage of established species that had invaded was 24%, a figure is 2.4 times higher than predicted by the tens rule.

Invasion rates were often significantly higher than predicted by the tens rule, but varied with the spatial scale at which they were estimated (Fig. 1). At the continental scale (L1), 25% (21-30%; 95% CI) of established plants have become invasive, while at the regional scale (L2), invasion rates were lower, although not significantly so, averaging 22% (16-27%; 95% CI). However, at the country/state scale (L3), invasion rates were significantly lower than the continental scale, averaging 17% (15-20%; 95% CI). At the sub-country scale (L4), invasion rates were significantly lower than the country/state scale, averaging 12% (9-14%; 95% CI), but data at this spatial resolution were rare (Fig. 1). Indeed, our dataset included data for all nine L1 continents, 48 of 52 L2 regions, 230 of 369 L3 countries, but only 68 of 315 L4 sub-country areas.

Invasion rates of non-native plants were significantly higher on islands than on mainlands (Fig. 2). This pattern was consistent across scales. At the global scale, invasion rates on islands averaged 19% (18-20%; 95% CI), while invasion rates on mainlands averaged 15% (14-16%; 95% CI). Similarly, at the country/state scale, when weighted by total number of established plants, invasion rates on islands averaged 18% (13-23%; 95% CI) and mainlands averaged 9% (8-11%; 95% CI).

**Fig. 2.**
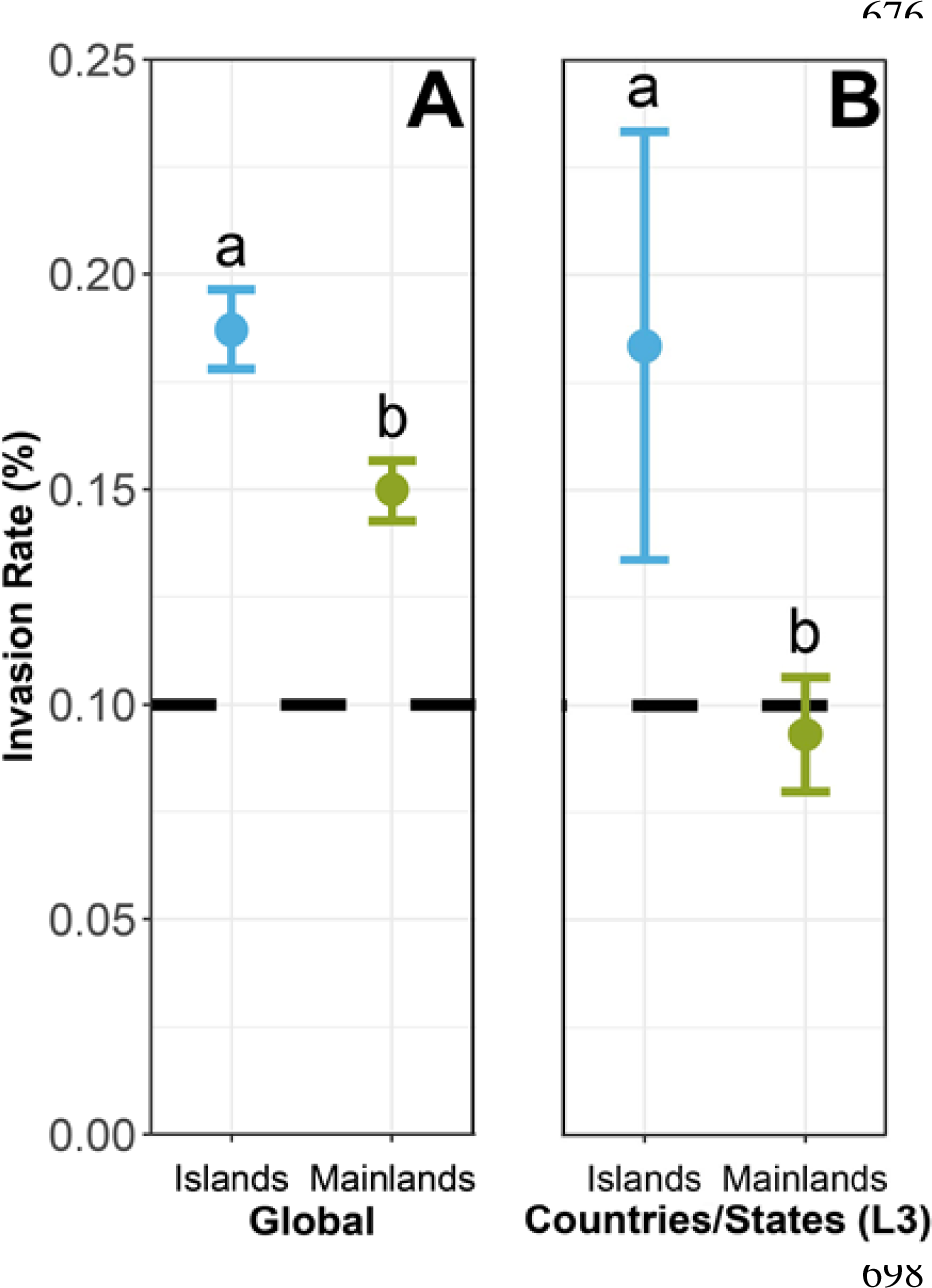
Islands have significantly higher invasion rates than mainlands. Invasion rates and 95% confidence intervals for islands and mainlands at the global scale (A) and the country/state (WGSRPD L3) scale (B). L3 averages are weighted by the total non-native species present in each of the regions used in the mean. The dashed line represents the tens rule. Lowercase letters (a, b) indicate significant differences (p-value ≤ 0.05) within panels. Test statistics, degrees of freedom, and p-values are provided in Table S3.

Invasion rates were significantly higher in the tropics than in arid, temperate, continental, or polar regions (Fig. 3). Overall, 25% (24-27%; 95% CI) of species that established in tropical regions became invasive, compared with 16% (15-17%; 95% CI) for arid regions, 14% (13-15%; 95% CI) for temperate regions, 8% (7-9%; 95% CI) for continental regions, and 10% (8-12%; 95% CI) for polar regions. Invasion rates for continental and polar climates were the most similar to the tens rule, but were significantly lower than all other climates. Invasion rates in tropical climates were significantly higher than all other climates, and were more than twice as high as estimated by the tens rule. These trends in the invasion rates of recipient climates were consistent regardless of the source climate(s) of the species (Fig. S1).

**Fig. 3.**
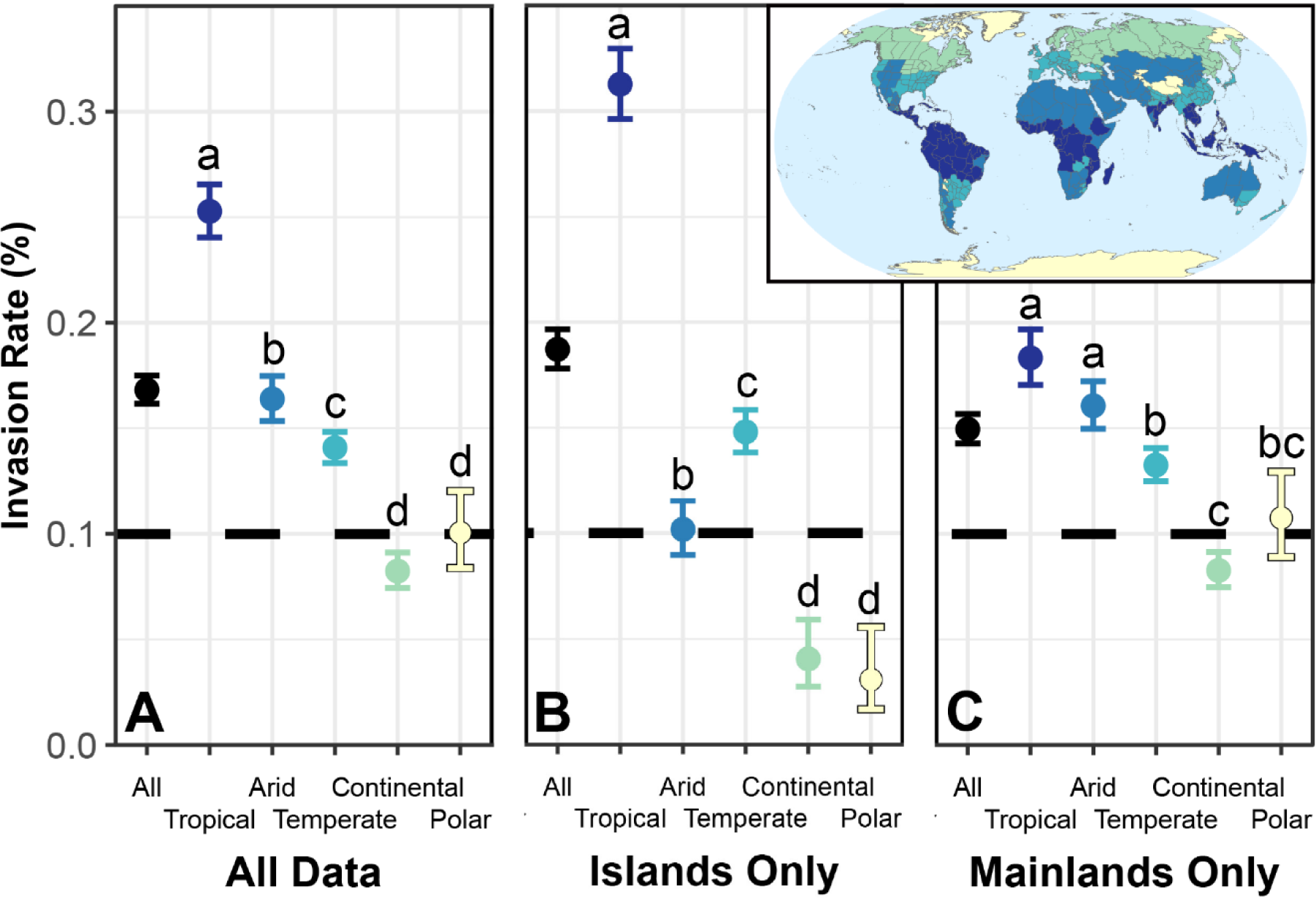
Tropical regions are particularly susceptible to invasion. Invasion rates and 95% confidence intervals within each of the 5 major Köppen climates for all regions with available data (A), only island regions (B), and only mainland regions (C). The dashed line represents the tens rule. Lowercase letters (a, b, c, d) indicate significant differences within panels (p-value ≤ 0.05). P-values are provided in Table S4.

We saw the same overall pattern of susceptibility varying by climate for both islands and mainlands (Fig. 3). However, there was a strong interaction between climate and island susceptibility (Fig. 3). The invasion rate on tropical islands was 31% (30-33%; 95% CI), which was significantly higher than islands within any other climate. Tropical islands also had significantly higher invasion rates than tropical mainlands.

## Discussion

### The tens rule does not represent global invasion rates

As originally presented by Williamson and Fitter (1996), the tens rule suggests that invasion risk is generally low, with only one in 100 introduced species successfully becoming invasive (Jarić and Cvijanović 2012). Jeschke and Pyšek (2018) suggest that the tens rule might be a “zombie idea”: a hypothesis that continues to be persistently communicated despite little empirical evidence. Our results categorically support this classification and reject the second transition within the tens rule. Considered globally, invasion rates for established plants are closer to 20%, which represents a doubling of invasion risk for the highly-impactful establishment-to-invasion transition.

Some existing estimates of invasion rates are consistent with our results, but the sample sizes and geographic scopes of these studies were extremely limited. For example, previous estimates of invasion rates in Great Britain (Williamson 1993, Williamson and Fitter 1996, Jeschke and Pyšek 2018) average 17.8%, and our estimate for temperate islands at the L3 scale is 13.8% (Table 1). Likewise, previous estimates for Australia average 20.1% (Scott and Panetta 1993, Virtue et al. 2004, Jeschke and Pyšek 2018), and our results project invasion rates for arid mainlands at the L2 scale to be 18.2% (Table 1). Importantly, however, our results also include invasion rates for these climates at other spatial scales, and we report percentages for continental, polar and tropical climates. Since the majority of the world’s biodiversity is found in tropical ecosystems (Raven 1988), and since invasion rates are scale-dependent, these previous estimates from only two climates at different spatial scales were unsatisfactory for drawing generalized conclusions. Indeed, Jeschke and Pyšek (2018) analyzed six studies whose results match our estimates, but they notably did not propose an alternative to the tens rule for plants due to the variability (individual studies ranged from 2% to 30%) and deficiency (only one study used a sample size greater than 300 plant species) of available data (Scott and Panetta 1993, Williamson 1993, Mills et al. 1993, Williamson and Fitter 1996, Virtue et al. 2004, Pyšek et al. 2012). In contrast, our data are comprehensive enough to extract global trends in invasion rates, and subsequently yield three novel conclusions: 1) for plants, 10% is not an appropriate estimate of global invasion rates, 2) invasion rates vary predictably with scale and geography, and 3) tropical ecosystems are especially vulnerable to invasion.

**Table 1.** Invasion rates by scale, climate, and islands/mainland status.

| Islands |  |  |  |  |  |
| --- | --- | --- | --- | --- | --- |
|  | Tropical | Arid | Temperate | Continental | Polar |
| **Regional (L2)** | 33.2 %<br>n<sub>REG</sub> = 8<br>n<sub>SPP</sub> = 2557 | 10.7 %<br>n<sub>REG</sub> = 1<br>n<sub>SPP</sub> = 954 | 26.0 %<br>n<sub>REG</sub> = 2<br>n<sub>SPP</sub> = 3018 | N/A<br>n<sub>REG</sub> = 0<br>n<sub>SPP</sub> = 0 | 10.6 %<br>n<sub>REG</sub> = 1<br>n<sub>SPP</sub> = 161 |
| **National (L3)** | 35.0 %<br>n<sub>REG</sub> = 39<br>n<sub>SPP</sub> = 2284 | 24.9 %<br>n<sub>REG</sub> = 4<br>n<sub>SPP</sub> = 370 | 13.8 %<br>n<sub>REG</sub> = 16<br>n<sub>SPP</sub> = 4800 | 4.5 %<br>n<sub>REG</sub> = 2<br>n<sub>SPP</sub> = 382 | 16.3 %<br>n<sub>REG</sub> = 42<br>n<sub>SPP</sub> = 2635 |
| **Sub-National (L4)** | 33.9 %<br>n<sub>REG</sub> = 43<br>n<sub>SPP</sub> = 1556 | 0 %<br>n<sub>REG</sub> = 1<br>n<sub>SPP</sub> = 1345 | 3.3 %<br>n<sub>REG</sub> = 14<br>n<sub>SPP</sub> = 2474 | 2.5 %<br>n<sub>REG</sub> = 3<br>n<sub>SPP</sub> = 436 | 0 %<br>n<sub>REG</sub> = 1<br>n<sub>SPP</sub> = 143 |
| Mainlands |  |  |  |  |  |
|  | Tropical | Arid | Temperate | Continental | Polar |
| **Regional (L2)** | 23.9 %<br>n<sub>REG</sub> = 9<br>n<sub>SPP</sub> = 2468 | 18.2 %<br>n<sub>REG</sub> = 14<br>n<sub>SPP</sub> = 5047 | 18.3 %<br>n<sub>REG</sub> = 7<br>n<sub>SPP</sub> = 4571 | 7.2 %<br>n<sub>REG</sub> = 8<br>n<sub>SPP</sub> = 3814 | 8.6 %<br>n<sub>REG</sub> = 2<br>n<sub>SPP</sub> = 336 |
| **National (L3)** | 18.6 %<br>n<sub>REG</sub> = 52<br>n<sub>SPP</sub> = 2804 | 15.0 %<br>n<sub>REG</sub> = 58<br>n<sub>SPP</sub> = 3829 | 13.3 %<br>n<sub>REG</sub> = 66<br>n<sub>SPP</sub> = 6369 | 8.6 %<br>n<sub>REG</sub> = 45<br>n<sub>SPP</sub> = 3561 | 7.2 %<br>n<sub>REG</sub> = 9<br>n<sub>SPP</sub> = 849 |
| **Sub-National (L4)** | 6.1 %<br>n<sub>REG</sub> = 43<br>n<sub>SPP</sub> = 1519 | 2.7 %<br>n<sub>REG</sub> = 56<br>n<sub>SPP</sub> = 2676 | 9.1 %<br>n<sub>REG</sub> = 90<br>n<sub>SPP</sub> = 3843 | 7.8 %<br>n<sub>REG</sub> = 11<br>n<sub>SPP</sub> = 1289 | 7.2 %<br>n<sub>REG</sub> = 5<br>n<sub>SPP</sub> = 531 |

n<sub>REG</sub> is the number of regions with each set of criteria. n<sub>SPP</sub> is the number of non-native plant species present across the regions.

Our study deliberately focuses on rates of *invasion* because they have previously been understudied and have significant consequences for policy and management. However, rates of *establishment* are also likely to be higher than 10%. For example, Jeschke and Pyšek identified eight studies (Williamson 1993, Williamson and Fitter 1996, Frenot et al. 2001, Křivánek and Pyšek 2006, Lambdon et al. 2008, Ööpik et al. 2008, Dawson et al. 2009, Pyšek et al. 2012) spanning 13 locations that measured the percentage of introduced plants that successfully established. The average establishment rate was 30% (22-38%; 95% CI), which is unsurprising, since the majority of plant introductions are deliberate, and many deliberately introduced plants are ornamentals (Virtue et al. 2004, Ööpik et al. 2008, Lehan et al. 2013). Ornamental plants are often pre-selected for tolerance of a variety of abiotic conditions, have high numbers of propagules, and benefit from repeated, widespread introduction and active cultivation by the people invested in their survival (Lockwood et al. 2005, Minton and Mack 2010, Briski et al. 2018). Williamson and Fitter (1996) described establishment rates of “crop plants” as an exception to the tens rule, writing “The tens rule does not apply when there has been selection to counteract it.” However, non-native plant cultivation (which nearly always involves some form of artificial selection) is so ubiquitous that considering this pathway as an exception ignores the primary mechanism of plant establishment (Mack and Erneberg 2002).

These updated estimates for establishment and invasion rates suggest global invasion risk is six times greater than previously estimated by the tens rule. In other words, if 30% of introduced species become established, and 20% of established species become invasive, then we should expect six percent of plants introduced to a new environment to become invasive. These rates likely represent underestimates, as lag phases between establishment and invasion are common in invasive plants (Aikio et al. 2010) and invasive plants are underreported in the Global South (Laginhas et al. 2022). Furthermore, new plant introductions continue to increase worldwide (Seebens et al. 2017), underscoring the need for this updated understanding of associated invasion risk.

The tens rule’s influence on existing invasive species policy is particularly concerning and emphasizes the importance of adopting more accurate estimates of invasion risk. For example, within the United States, many invasive species management plans and risk assessments at both the state (L3) and federal (L2) levels have directly referenced the tens rule (Daehler and Carino 2000, Northwest Natural Resource Group 2003, Carlson et al. 2008, Koop et al. 2012, Minnesota Noxious Weed Advisory Committee 2020). In these reports, the tens rule is often described as a precise estimate of invasion rates, or as an upper limit for the percent of non-native species that could pose negative impacts. This interpretation represents an unfortunate oversimplification of the wording used by Williamson and Fitter (1996). Moreover, practitioners have frequently referred to the tens rule in contexts outside of manuscripts and reports, which suggests that current management decisions and invasive species policies are influenced by the tens rule to a greater extent than is explicitly documented (Davis and Hoff 2016, Maryland Department of Natural Resources 2023, Native Plant Trust 2023). Given the higher invasion rates reported here in comparison to the existing use of the tens rule in management scenarios, it is likely that these systems have consistently overestimated our ability to correctly predict and prevent the introduction of invasive plants, and therefore also have consistently underestimated the potential consequences of nonnative species. In the future, managers should consider the spatial scale, island status, and climate of their targeted ecosystems to gain a more nuanced and precise understanding of risk (Table 1).

### Scale dependence of invasion rates

Invasion rates show a clear scale dependence, supporting the hypothesis that invasion rates increase when study areas are large. This is likely because larger areas encompass a greater diversity of habitats to invade such that widely established species might cause negative impacts in a single location, even if not consistently invasive throughout the broader area (Jeschke and Pyšek 2018) (Fig. 1). Since this pattern has been unaddressed in existing literature, it has likely contributed to the varied support for the tens rule provided by different studies (e.g., Williamson and Fitter 1996, Virtue et al. 2004, Pyšek et al. 2012, Jeschke and Pyšek 2018). Our results clearly demonstrate the effect of spatial scale, as invasion rates are highest at the coarsest scale and decrease at each subsequently finer scale. Although we calculated invasion rates for four spatial scales, in practice, regulations and risk assessments for invasive plants are most common at the regional (L2) or country/state (L3) scales (Salva 2023). At these scales, invasion rates of 20% are a better rule of thumb for evaluating invasion risk and informing regulation of introduced plants.

### Island susceptibility

Our results support previous findings that islands are more susceptible to invasion (Elton 1958, Simberloff 1995, Lonsdale 1999) (Fig. 2). Islands may be more susceptible because biotic resistance is weaker due to evolutionary naivete (Simberloff 1995, Rejmánek 1996). Additionally, the theory of island biogeography suggests that resource rich islands contain fewer species than they could potentially support, leading to more opportunities for invasion when compared to mainland areas with similar abiotic conditions (MacArthur and Wilson 1967, Jeschke et al. 2018). Significantly higher invasion rates on tropical islands suggest that invasive plants are able to take advantage of underutilized island resources, with one third of established plants on tropical islands identified as invasive (Fig. 3).

The notion that island susceptibility is driven by the interaction of resource availability and biotic factors is further supported by the small differences in invasion rates we observed between islands and mainlands in polar regions. As hypothesized by Darwin (1859) and demonstrated more recently (Paquette and Hargreaves 2021), the relative importance of abiotic factors (i.e. resource availability) compared to biotic factors (i.e. competition) should increase in higher latitudes. Therefore, although both polar and tropical islands likely have reduced competition due to fewer native species, the overall lack of available resources in polar climates appears to decrease invasion rates similarly on both islands and mainlands, resulting in decreased support for island susceptibility at higher latitudes (Fig. 3).

### Tropical susceptibility

Tropical areas have high native species richness and have accordingly been hypothesized to have higher resistance to invasions (Rejmánek 1996, Chong et al. 2021) due to biotic resistance. While several studies have found fewer plant invasions in the tropics than other climates (e.g. Rejmánek 1996, Lonsdale 1999), tropical invasions are also chronically understudied. Chong et al. (2021) recently hypothesized that invasion rates in the tropics may be comparable to temperate regions since low numbers of invasive plants in the tropics may be an artifact of fewer species introductions, fewer anthropogenic disturbances and/or sampling biases. However, our results suggest that tropical regions are in fact significantly more susceptible to invasion when compared to arid, temperate, continental, and polar regions (Fig. 3). This novel insight emerging from our analysis has wide-ranging and significant implications for many ecological and evolutionary hypotheses about the nature of tropical communities. Even when we only compared mainland areas (controlling for the influence of island susceptibility), we still found that the tropics had the highest invasion rates of any climate. Previous work has shown that areas with higher resources are more susceptible to invasion, particularly in the presence of disturbance or absence of natural enemies (Davis et al. 2000, Blumenthal 2006), but these studies have largely been conducted within continental areas or across habitats within regions. That a similar pattern might be present at global scales is striking and runs counter to many predictions based on evidence of increased intensity of antagonistic biotic interactions in the tropics. In contrast to tropical regions, polar and continental climates had the lowest invasion rates. Large annual fluctuations in climate conditions may create strong abiotic limitations for species invasion in these regions (Janzen 1967, Zefferman et al. 2015).

Thus, at regional scales, our results suggest that greater abiotic resource availability might counteract any biotic resistance conferred from higher species richness in the tropics (Davis et al. 2000, Blumenthal 2006). While biotic resistance generally holds at local scales (Levine et al. 2004), regional invasion rates mirror abiotic resource gradients. Currently, invasive plant richness in the tropics (relative to native diversity) remains low (Laginhas et al. 2022), but our results highlight an elevated risk of future invasions, particularly in light of ongoing introductions (Seebens et al. 2017) and increasing disturbance (Myers 1994).

### Future management

Future management of invasive plant species should be adjusted to better reflect the invasion rates described here. For instance, one of the most common invasive species management strategies is “early detection and rapid response” (EDRR), which seeks to eradicate new invasive populations before they spread (Reaser et al. 2020). Managers of locations that are more vulnerable to invasion (e.g. tropical islands), should invest a greater proportion of resources to EDRR. Since many newly-established plant species are likely to cause harm in these locations, eradicating emerging populations while they are in the early stages of invasion is paramount. In contrast, where invasion rates are low (e.g. mainlands with continental climates), management investment should instead go towards controlling existing invasions. In addition to informing EDRR, invasion rates are also critical for informing risk assessment, quarantine, and prevention policies at national borders. Our results show that tropical countries throughout much of the Global South are likely to experience high rates of invasion. Unfortunately, many of these countries lack national-scale weed risk assessments (Salva 2023) and are therefore in dire need of improved policies to defend against invasive species introduction.

## Conclusion

In their original tens rule article, Williamson and Fitter (1996) acknowledged that the tens rule was not meant to be a precise indicator of invasion rates, instead noting that it was a “crude” estimate, which may actually fluctuate between 5 and 20%. Nevertheless, their 10% estimate has been widely and pervasively repeated in scientific reports and popular media for almost three decades (see for example Carlson et al. 2008, Davis and Hoff 2016, United States Environmental Protection Agency 2016, Cary Institute of Ecosystem Studies 2020, Minnesota Noxious Weed Advisory Committee 2020, Shah 2020). Given that invasive plants are responsible for billions of dollars in annual damages and declines of native species (Pimentel et al. 2005, Vilà et al. 2011), it is imperative that we accurately calculate and communicate empirically-grounded, globally-based invasion risk estimates. Moving forward, researchers and practitioners should use the context-specific estimates of invasion rates provided here, especially for high-value, high-diversity tropical regions that are of significant global conservation concern. Legislation and management practices should be updated accordingly to reflect this heightened likelihood of invasion, especially in tropical climates and island habitats. Continuing to use the tens rule as a global rule of thumb for invasion risk would sustain underestimates of ecological and economic consequences of non-native plants worldwide, as well as perpetuate the harmful misconception that an insignificant number of established species become invasive.

## Acknowledgments

We thank Michael Gonthier, David Matevosian, Julia Mazzuchi, Fatimah Rashid, and Michael Wilkinson for their assistance with data collection. We thank Mark van Kleunen and the GloNAF team for their willingness to share data. We also thank the members of the Spatial Ecology Lab at UMass, Daniel Buonaiuto, Laura Figueroa, Jesse Bellemare, Graziella DiRenzo, and Matthew Fertakos for their feedback on analyses, interpretation, phrasing, and figures.

## Funding

Lotta Crabtree Trust (WGP), U.S. Geological Survey Northeast Climate Adaptation Science Center graduate fellowship, grant G19AC00091 (WGP)

## Author contributions

Conceptualization: WGP, BAB. Data Collection: WGP. Methods: WGP, BAB. Analysis and Visualization: WGP. Funding Acquisition: WGP, BAB. Writing: WGP, BAB.

## Conflict of interest statement

The authors declare that they have no conflicts of interest to report.

## Notes

https://github.com/wpfadenhauer/Global-Invasion-Rates

### Competing Interest Statement

The authors have declared no competing interest.

