## Supplementary Materials for "Quantifying vulnerability to plant invasion across global ecosystems"

*Ecological Applications*

William G. Pfadenhauer*, Bethany A. Bradley

**This PDF file includes:**

Figs. S1 to S2

Table S1 to S4

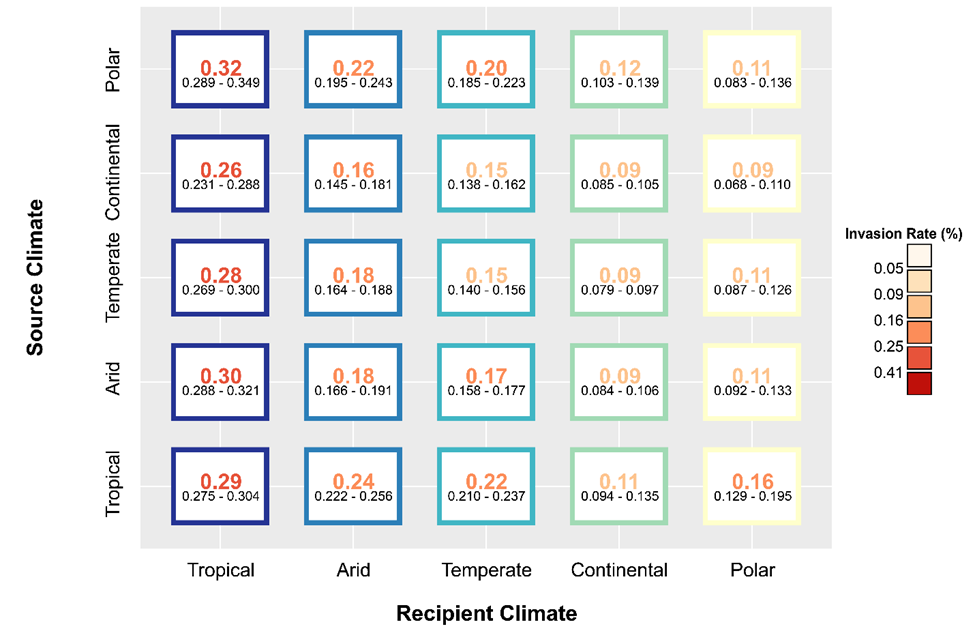

Fig. S1.

Invasion rates broken down by source and recipient climates. The number in the center of each tile is the invasion rate, which is color-coded to match Figure 1. The border color of each tile corresponds to the recipient climate. Smaller, black numbers represent 95% confidence intervals.

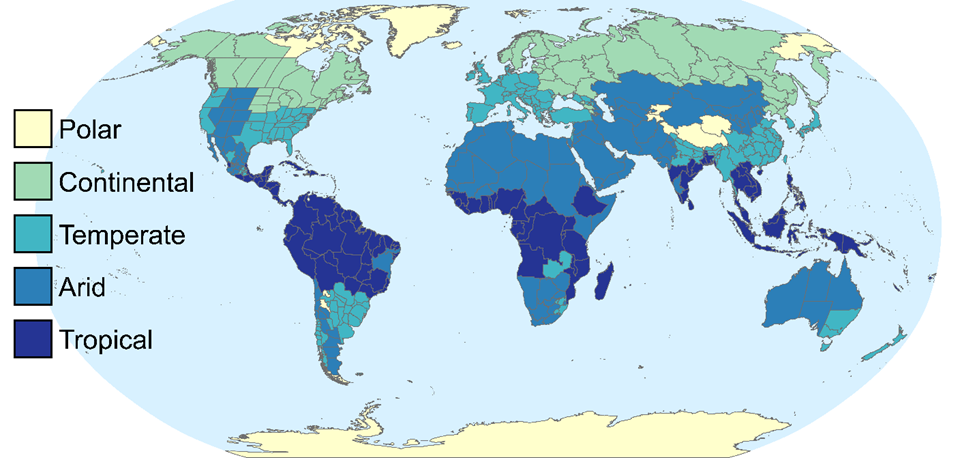

Fig. S2.

The Köppen climate that is most common within each of the WGSRPD L3 / L4 regions (whichever level is the finest scale available in each location).

Table S1.

Sources and methods used to collect data for the native, established, and invaded distributions for each of the 12,425 candidate species. Referenced cited here are in the reference list for the main text.

| **Data** | **Source** | **Method** |
| --- | --- | --- |
| Native distributions | CABI Invasive Species Compendium  (CAB International 2022) | Webscraped |
|  | Plants of the World Online  (Royal Botanic Gardens, Kew 2022) | Webscraped |
| Established distributions | Global Naturalized Alien Flora  (van Kleunen et al. 2019) | Downloaded |
| Invasive distributions | GRIIS  (Global Register of Introduced and Invasive Species 2022) | Downloaded |
|  | CABI Invasive Species Compendium  (CAB International 2022) | Webscraped |

Table S2.

T-values, degrees of freedom (df) and p-values for Figure 1.

| **Data** | **Comparison** | **t-value** | **df** | **p-value** |
| --- | --- | --- | --- | --- |
| L1 invasion rates | 10% | 7.7849 | 8 | 5.311e-05 |
| L2 invasion rates | 10% | 4.4567 | 50 | 4.705e-05 |
| L3 invasion rates | 10% | 5.7985 | 250 | 2.012e-08 |
| L4 invasion rates | 10% | 1.1414 | 129 | 0.2558 |
| L1 invasion rates | L2 invasion rates | 1.0504 | 41.64 | 0.2996 |
| L1 invasion rates | L3 invasion rates | 3.4081 | 15.91 | 0.003622 |
| L1 invasion rates | L4 invasion rates | 5.6859 | 17.23 | 2.239e-05 |
| L2 invasion rates | L3 invasion rates | 1.529 | 74.52 | 0.1305 |
| L2 invasion rates | L4 invasion rates | 3.4133 | 79.13 | 0.001014 |
| L3 invasion rates | L4 invasion rates | 3.0477 | 317.4 | 0.002499 |

Table S3.

Test statistics, degrees of freedom (df) and p-values for Figure 2. The test statistic for row 1 (panel A) is a X-squared value (proportion test), and the test statistic for row 2 (panel B) is a t-value (weighted t-test).

| **Data** | **Comparison** | **Statistic** | **df** | **p-value** |
| --- | --- | --- | --- | --- |
| Global islands | Global mainlands | 41.746 | 1 | 5.198e-11 |
| L3 islands | L3 mainlands | 3.547 | 57.71 | 3.913e-04 |

Table S4.

P-values for Figure 3.

| **Panel** | **Data** | **Comparison** | **p-value** |
| --- | --- | --- | --- |
| A | Tropical | Arid | < 2e-16 |
| A | Tropical | Temperate | < 2e-16 |
| A | Tropical | Continental | < 2e-16 |
| A | Tropical | Polar | < 2e-16 |
| A | Arid | Temperate | 0.0039 |
| A | Arid | Continental | < 2e-16 |
| A | Arid | Polar | 3.4e-06 |
| A | Temperate | Continental | < 2e-16 |
| A | Temperate | Polar | 0.0042 |
| A | Continental | Polar | 0.666 |
| B | Tropical | Arid | < 2e-16 |
| B | Tropical | Temperate | < 2e-16 |
| B | Tropical | Continental | < 2e-16 |
| B | Tropical | Polar | < 2e-16 |
| B | Arid | Temperate | 2.3e-06 |
| B | Arid | Continental | 3.3e-05 |
| B | Arid | Polar | 0.00058 |
| B | Temperate | Continental | 3.1e-12 |
| B | Temperate | Polar | 5.9e-08 |
| B | Continental | Polar | 1.0 |
| C | Tropical | Arid | 0.10369 |
| C | Tropical | Temperate | 1.1e-10 |
| C | Tropical | Continental | < 2e-16 |
| C | Tropical | Polar | 8.1e-07 |
| C | Arid | Temperate | 0.00043 |
| C | Arid | Continental | < 2e-16 |
| C | Arid | Polar | 0.00068 |
| C | Temperate | Continental | 8.5e-15 |
| C | Temperate | Polar | 0.39641 |
| C | Continental | Polar | 0.18662 |

van Kleunen, M., P. Pyšek, W. Dawson, F. Essl, H. Kreft, J. Pergl, P. Weigelt, A. Stein, S. Dullinger, C. König, B. Lenzner, N. Maurel, D. Moser, H. Seebens, J. Kartesz, M. Nishino, A. Aleksanyan, M. Ansong, L. A. Antonova, J. F. Barcelona, S. W. Breckle, G. Brundu, F. J. Cabezas, D. Cárdenas, J. Cárdenas‐Toro, N. Castaño, E. Chacón, C. Chatelain, B. Conn, M. de S. Dechoum, J.-M. Dufour‐Dror, A. L. Ebel, E. Figueiredo, O. Fragman‐Sapir, N. Fuentes, Q. J. Groom, L. Henderson, Inderjit, N. Jogan, P. Krestov, A. Kupriyanov, S. Masciadri, J. Meerman, O. Morozova, D. Nickrent, A. Nowak, A. Patzelt, P. B. Pelser, W. Shu, J. Thomas, A. Uludag, M. Velayos, A. Verkhosina, J. L. Villaseñor, E. Weber, J. J. Wieringa, A. Yazlık, A. Zeddam, E. Zykova, and M. Winter. 2019. The Global Naturalized Alien Flora (GloNAF) database. Ecology 100:e02542.

Royal Botanic Gardens, Kew. 2022. Plants of the World Online. https://powo.science.kew.org/.
